# The ability of transcription factors to differentially regulate gene expression is a crucial component of the mechanism underlying inversion, a frequently observed genetic interaction pattern

**DOI:** 10.1101/449520

**Authors:** Saman Amini, Annika Jacobsen, Olga Ivanova, Philip Lijnzaad, Jaap Heringa, Frank C. P. Holstege, K. Anton Feenstra, Patrick Kemmeren

**Author notes:** These authors contributed equally to this work (Saman Amini, Annika Jacobsen).

## Abstract

Genetic interactions, a phenomenon whereby combinations of mutations lead to unexpected effects, reflect how cellular processes are wired and play an important role in complex genetic diseases. Understanding the molecular basis of genetic interactions is crucial for deciphering pathway organization as well as understanding the relationship between genetic variation and disease. Several putative molecular mechanisms have been linked to different genetic interaction types. However, differences in genetic interaction patterns and their underlying mechanisms have not yet been compared systematically between different functional gene classes. Here, differences in the occurrence and types of genetic interactions are compared for two classes, gene-specific transcription factors (GSTFs) and signaling genes (kinases and phosphatases). Genome-wide gene expression data for 63 single and double deletion mutants in baker’s yeast reveals that the two most common genetic interaction patterns are buffering and inversion. Buffering is typically associated with redundancy and is well understood. In inversion, genes show opposite behavior in the double mutant compared to the corresponding single mutants. The underlying mechanism is poorly understood. Although both classes show buffering and inversion patterns, the prevalence of inversion is much stronger in GSTFs. To decipher potential mechanisms, a Petri Net modeling approach was employed, where genes are represented as nodes and relationships between genes as edges. This allowed over 9 million possible three and four node models to be exhaustively enumerated. The models show that a quantitative difference in interaction strength is a strict requirement for obtaining inversion. In addition, this difference is frequently accompanied with a second gene that shows buffering. Taken together, these results provide a mechanistic explanation for inversion. Furthermore, the ability of transcription factors to differentially regulate expression of their targets provides a likely explanation why inversion is more prevalent for GSTFs compared to kinases and phosphatases.

**Author Summary:** The relationship between genotype and phenotype is one of the major challenges in biology. While many previous studies have identified genes involved in complex genetic diseases, there is still a gap between genotype and phenotype. One of the difficulties in filling this gap has been attributed to genetic interactions. Large-scale studies have revealed that genetic interactions are widespread in model organisms such as baker’s yeast. Several molecular mechanisms have been proposed for different genetic interaction types. However, differences in occurrence and underlying molecular mechanism of genetic interactions have not yet been compared between gene classes of different function. Here, we compared genetic interaction patterns identified using gene expression profiling for two classes of genes: gene specific transcription factors and signaling related genes. We modelled all possible molecular networks to unravel putative molecular differences underlying different genetic interaction patterns. Our study proposes a new mechanistic explanation for a certain genetic interaction pattern that is more strongly associated with transcription factors compared to signaling related genes. Overall, our findings and the computational methodologies implemented here can be valuable for understanding the molecular mechanisms underlying genetic interactions.

## Introduction

Understanding the relationship between genotype and phenotype of an organism is a major challenge [1,2]. One of the difficulties for unravelling genotype-phenotype relationship has been genetic interactions, when combinations of mutations lead to phenotypic effects that are unexpected based on the phenotypes of the individual mutations [3–5]. Large-scale analyses of single and double deletion mutants have revealed that genetic interactions are pervasive in many model organisms [6–11]. Recently, efforts have been initiated to investigate genetic interactions in human cell lines too, using large-scale RNA interference and Crispr-Cas9 knock downs [12–15]. Our understanding of the molecular mechanisms that underlie genetic interactions lags behind our ability to detect genetic interactions. Understanding the molecular basis of genetic interactions and their interplay with cellular processes is important for unraveling how different processes are connected [16–18], to what degree genetic interactions shape pathway architecture [6], as well as for understanding the role genetic interactions play in human disease [5,19].

One of the phenotypes that is frequently used to investigate genetic interactions is cell growth [6,20–28]. Based on this phenotype, genetic interactions can be broadly subdivided in two types, negative genetic interactions where the double mutant is growing slower than expected given the growth rate of the single deletion mutants, and positive genetic interactions where the double mutant is growing faster than expected [3]. Negative genetic interactions have frequently been associated with a redundancy relationship between two functionally related genes [29]. The redundancy mechanisms by which two genes can compensate for each other’s loss has been linked with close paralog genes or redundant pathways [30,31]. Positive genetic interactions have been associated with genes participating in the same protein complex or pathway [32]. There are however many exceptions to these rules and it also has become clear that there are many other potential mechanisms underlying these genetic interactions [3,18].

Another phenotype that has been less frequently used for investigating genetic interactions is gene expression [16,17,33–36]. Expression-based genetic interaction profiling provides detailed information at the molecular level which is beneficial for unraveling mechanisms of genetic interactions [16,17,33–36]. Unlike growth-based profiling, which gives a subdivision into either positive or negative interactions, expression-based genetic interaction profiling provides further subdivision into more specific genetic interaction patterns including buffering, quantitative buffering, suppression, quantitative suppression, masking and inversion [17]. A more detailed sub classification that includes information on expression of downstream genes, can also contribute to understanding the mechanisms by which two genes interact [16,17,37].

To provide mechanistic insights into biological networks, Boolean modeling has been used successfully [38,39]. It has also been applied to unravel regulatory networks underlying genetic interaction patterns between kinases and phosphatases [16]. Due to their intrinsically simple nature, such Boolean network models allow exhaustive enumeration of network topologies. The outcomes of these models can then be easily compared to the patterns observed in experimental data. Boolean operators however, are limited to on and off values and cannot easily accommodate quantitative measurements, which limits the types of genetic interaction patterns that can be investigated using this approach. Unravelling the regulatory network underlying genetic interaction patterns would potentially benefit from application of modeling approaches that allow some degree of quantitativeness to be introduced while still being computationally feasible to exhaustively explore all potential models. In this way, Petri nets may be considered an extension of Boolean modeling that provides more flexibility, in particular by choosing different network edge strengths, without the need to incorporate detailed prior quantitative knowledge [40–44]. Petri net modeling would therefore allow investigation of all possible genetic interaction patterns in an exhaustive and semi-quantitative manner.

It is evident that genetic interactions are widespread in Saccharomyces cerevisiae [6] as well as other organisms [7,8]. Nevertheless, extensive characterization of the molecular mechanisms underlying genetic interactions, as well as a comparison of the molecular mechanisms underlying genetic interactions between different functional classes have, as yet, not been performed. Here, two functional classes, gene specific transcription factors (GSTFs) and signaling related genes (kinases and phosphatases) have been compared with regard to negative genetic interaction patterns and the possible underlying molecular mechanisms. This revealed that the two most common genetic interaction patterns are buffering and inversion. The prevalence of inversion however, is much stronger in GSTFs. The underlying mechanism of inversion, whereby genes show opposite behavior in the double mutant compared to the corresponding single mutants, is poorly understood. Exhaustive enumeration of network topologies using Petri net modelling reveals that the minimum requirement for observing inversion is having a quantitative difference in interaction strength (edge weight) from the two upstream transcription factors to a shared downstream gene. In addition, this quantitative edge difference is frequently accompanied by an intermediate node, that displays a buffering pattern. The proposed model provides a mechanistic explanation for inversion, thereby further aiding a better understanding of genetic interactions. GSTFs, more so than kinases/phosphatases, can modulate or fine-tune the activation levels of their target genes, which suggests quantitative differences in regulating downstream target genes are important for the functioning of GSTFs. This is consistent with the fact that inversion occurs more often between GSTFs than between signaling genes, as well as our observation that quantitative edge differences are required for inversion to occur and provides a likely explanation why inversion is more prevalent for transcription factors.

## Results

### A single dataset to compare mechanisms of genetic interactions between gene-specific transcription factors and kinases/phosphatases

To investigate potential differences in mechanisms of genetic interactions between groups of genes with a different function, data from two previously published datasets were combined [16,17]. The first dataset includes genome-wide gene expression measurements of 154 single and double gene-specific transcription factor (GSTF) deletion mutants [17]. The second dataset contains genome-wide gene expression measurements of 54 single and double kinase/phosphatase (K/P) deletion mutants [16]. These studies applied different criteria to select for interacting pairs. Whereas the GSTF dataset includes both positive and negative genetic interactions, the kinase/phosphatase dataset was restricted to negative genetic interactions only. To avoid potential biases, the selection criteria of the kinase/phosphatase dataset [16] were adopted and applied to both datasets. In short, selection was based on pairs having a significant growth-based negative genetic interaction score (p < 0.05, Methods) to include redundancy relationships that influence fitness. In addition, for a given double mutant, at least one of the corresponding single mutants has an expression profile similar to wildtype (WT) (eight or more transcripts changing significantly (p < 0.05, fold-change > 1.7)) to ensure that genetic interactions such as redundancy are considered. These selection criteria yield a uniform dataset consisting of 11 GSTF double mutants and 15 kinase/phosphatase double mutants as well as their respective single mutants (63 single and double mutants in total; S1 Table).

### Genetic interaction profiles indicate a large degree of buffering

Genetic interactions can be investigated in different ways. Here, both growth as well as genome-wide gene expression is used to compare genetic interactions between GSTFs and kinases/phosphatases, as described before [17]. In short, a growth-based genetic interaction score *ε_growth,XY_* between two genes X and Y is obtained by comparing the observed fitness for double mutant W*_xΔyΔ_* to the fitness that is expected based on both single mutants W*_xΔ ·_* W*_yΔ_* (ε*_growth,XY_* = W*_xΔyΔ_* - W*_xΔ_* · W*_yΔ_*)[45]. A gene expression-based genetic interaction score between two genes X and Y is calculated in two consecutive steps [17]. First, the effect of a genetic interaction between two genes X and Y on any downstream gene *i* is calculated as the deviation between the expression change observed in the double mutant M*_i,xΔyΔ_* and the expected expression change based on the corresponding single mutants M_*i,xΔ*_ + M_*i,yΔ*_ (ε_*txpn_iXY*_ = |M_*i,xΔyΔ*_ - (M_*i,xΔ*_+ M_*i,yΔ*_)|). The overall genetic interaction score between gene X and Y is then obtained by counting the total number of genes for which ε_*txpn_i,XY*_ is greater than 1.5 [17]. Gene expression changes from single and double mutants were subsequently grouped into the six genetic interaction patterns, buffering, suppression, quantitative buffering, quantitative suppression, masking and inversion, as previously described (Fig 1A) [17]. When investigating the genetic interaction profiles of GSTFs (Fig 1B) as well as kinases/phosphatases (Fig 1C), it is clear that buffering is prevalent in many of the larger genetic interaction profiles, but the degree of buffering differs for the smaller genetic interaction profiles.

**Fig 1.**
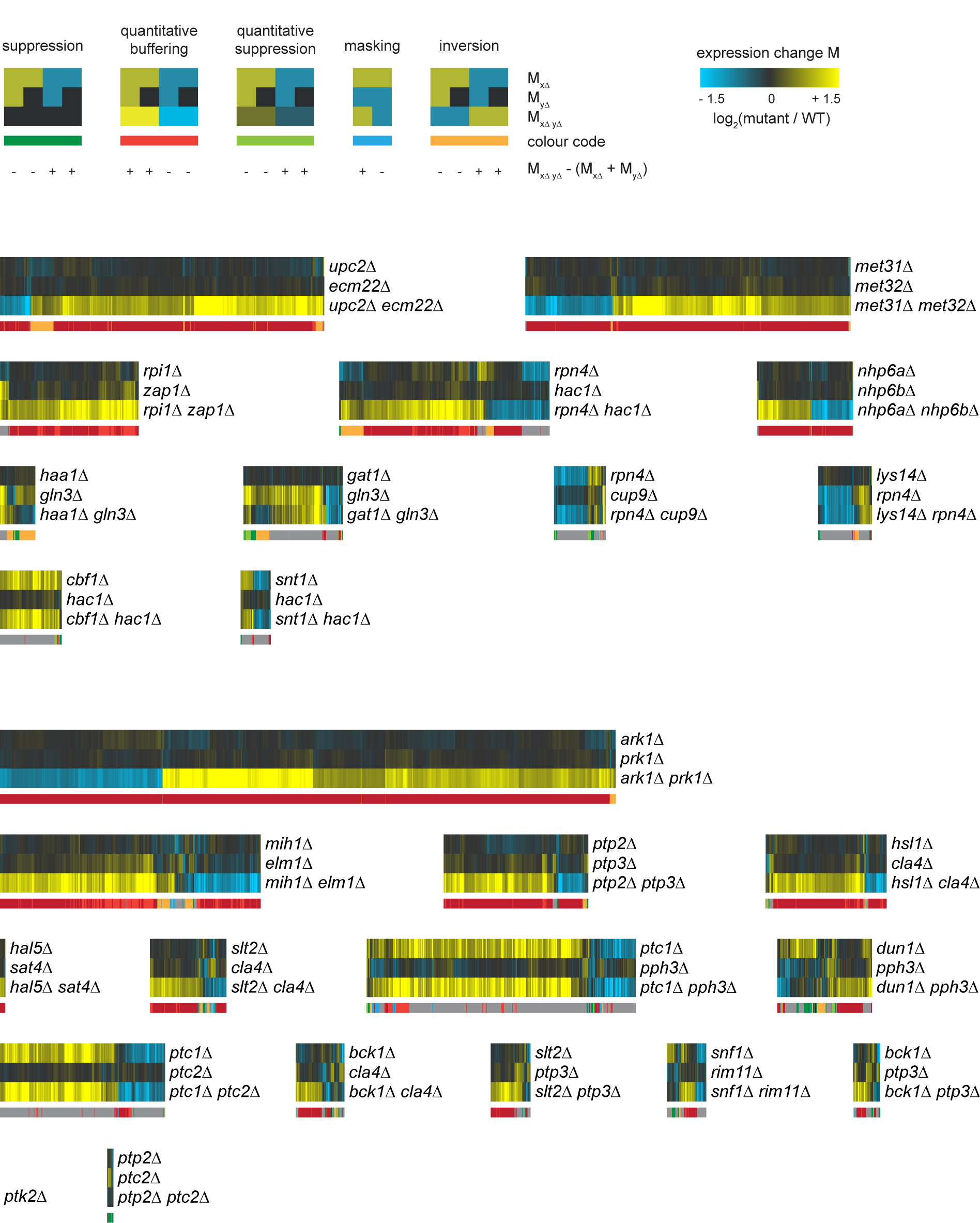
Genetic interaction profiles of GSTF and kinase/phosphatase pairs. **(A)** Cartoon depicting expression changes in single and double mutants with different genetic interaction patterns color coded underneath. At the bottom, the direction of expression differences between the observed expression change (M_*xΔyΔ*_) and expected (M_*xΔ*_+M_*yΔ*_) is stated. Color scale from yellow for an increase in expression levels compared to WT (p ≤ 0.01, log_2_(FC) > 0), black for unchanged expression (*p* > 0.01) and blue for a decrease in expression levels compared to WT (p ≤ 0.01, log_2_(FC) < 0). (B) Expression changes compared to WT (horizontal) in GSTF single and double mutants (vertical). Different colors underneath the gene expression profiles represent different genetic interaction patterns as indicated in A. Gray depicts gene expression changes not part of a genetic interaction pattern. Pairs are sorted based on the number of genetic interaction effects, increasing from bottom to top. (C) Expression changes compared to WT (horizontal) in kinase and phosphatase single and double mutants (vertical). Layout and ordering as in B.

### Removal of a slow growth associated expression signature for improved identification of direct effects

Hierarchical clustering was applied to group pairs with similar genetic interaction patterns (S1 Fig), thereby disregarding the identity of individual downstream genes. From this clustering, it is clear that there is no distinct separation between pairs consisting of GSTFs and kinases/phosphatases. Instead, most pairs are characterized by large buffering effects, grouped together in a single large cluster (S1A Fig, red branch labeled as 1). This is not surprising, since all pairs are selected for having a significant growth-based negative genetic interaction score. This in turn is based on double mutants growing slower than expected based on the single mutants. Slow growing strains are known to display a common gene expression signature [46,47]. This slow growth gene expression signature is caused by a change in the distribution of cells over different cell cycle phases [48]. To facilitate investigating mechanisms of genetic interactions, such effects are better disregarded. As described previously [48], the dataset was transformed by removing the slow growth signature (Methods). Removing the slow growth signature and thereby reducing effects due to a cell cycle population shift improves identification of direct target genes of GSTF pairs (S2 Fig) as shown before for individual GSTFs [48].

### Discerning potential mechanisms with slow growth corrected genetic interaction profiles

Hierarchical clustering of the slow growth corrected genetic interaction profiles was then applied to unravel potential differences in observed genetic interactions patterns between GSTFs and K/P (Fig 2A-C). Three striking differences emerge when comparing this clustering with the clustering of the original, untransformed data (S1 Fig). First, pairs are grouped into four distinct clusters, whereas previously, most were grouped into a single large cluster. Second, a cluster of predominantly kinase/phosphatase pairs emerges (Fig 2A, green branch, labeled as 1). These contain mixtures of different genetic interaction patterns, corresponding to ‘mixed epistasis’ [16]. Third, a smaller cluster dominated by buffering appears (Fig 2A, red branch, labeled as 2). This cluster also has strong growth-based negative genetic interaction scores (Fig 2C), which is known to be associated with redundancy. The ‘buffering’ cluster, with its strong growth-based negative interactions, mostly consists of pairs with a high sequence identity (average 43.7%) compared to the others (average 21%). These include Nhp6a-Nhp6b, Met31-Met32, Ecm22-Upc2 and Ark1-Prk1, for all of which redundancy relationships have been described previously [49–52]. The high sequence identity here indicates a homology-based redundancy, in which both genes can perform the same function [30,31,53,54]. The only exception here, is the kinase/phosphatase pair Elm1-Mih1. This pair may be explained through pathway-based redundancy where two parallel pathways can compensate for each other’s function [55]. Elm1 is a serine/threonine kinase, and Mih1 a tyrosine phosphatase, which are both involved in cell cycle control (S3 Fig, left panel) [56,57]. Mih1 directly regulates the cyclin-dependent kinase Cdc28, a master regulator of the G2/M transition [57]. Elm1, on the other hand, indirectly regulates Cdc28 activity by promoting Swe1 degradation through the recruitment of Hsl1 [58,59]. The timing of entry into mitosis is controlled by balancing the opposing activities of Swe1 and Mih1 on Cdc28, and both Swe1 and Mih1 are key in the checkpoint mediated G2 arrest [60,61]. Deletion of Elm1 does not result in many gene expression changes (Fig 1C) which can be explained through compensatory activity of Mih1 (S3 Fig, middle panel). Downregulation of Mih1 activity has also been suggested before as an effective mechanism to counter stabilization of Swe1, as neither stabilization of Swe1 or elimination of Mih1 in itself is sufficient to promote G2 delay, but simultaneous stabilization of Swe1 and elimination of Mih1 does cause G2 arrest [59]. Simultaneous deletion of Elm1 and Mih1 leads to higher levels of inactive Cdc28 causing a G2 delay and stress (S3 Fig, right panel) [59]. All pairs within this cluster can therefore be associated with a redundancy mechanism. Taken together, these results suggest that the clustering of the slow growth corrected genetic interaction profiles is able to discern potential differences in mechanisms. Even though most pairs in the four clusters (Fig 2A) show negative genetic interactions (Fig 2C), different mechanisms are likely underlying each individual cluster.

**Fig 2.**
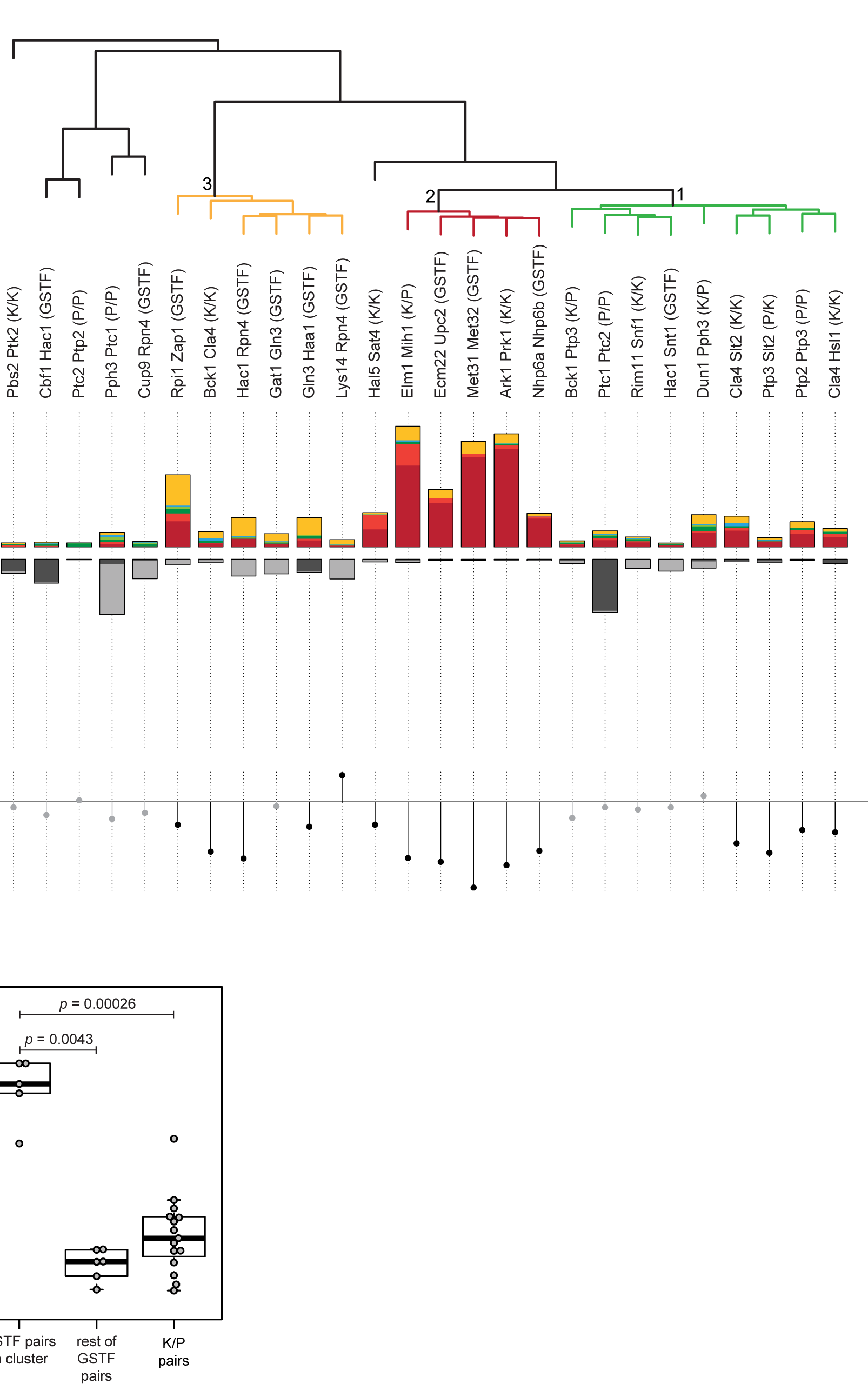
Hierarchical clustering of slow growth corrected genetic interaction profiles is better suited to discern underlying mechanisms. (A) Hierarchical clustering of all pairs according to their genetic interaction effects after slow growth correction. Average linkage clustering was applied to group pairs with similar genetic interaction patterns. The number of occurrences for each genetic interaction pattern (Fig 1A) was used and the identity of individual genes was disregarded. Similarity between pairs was calculated using cosine correlation. Branch depicted in red, label 2, indicates pairs that are dominated by buffering. Branch depicted in orange, label 3, indicates pairs dominated by inversion. Branch depicted in green, label 1, indicates pairs explained by mixed epistasis. The number of genetic interaction effects underlying the clustering are shown as bar plots below the dendrogram (colors as in Fig 1A). (B) Number of genes showing no genetic interaction pattern but significantly changing in one of the mutants compared to WT (p ≤ 0.01, FC > 1.5). Dark gray for the first named gene, light gray for the second named gene. (C) Growth-based genetic interaction scores depicted by solid circles. Significant genetic interaction scores are shown in black, gray otherwise. Ordering of pairs is the same as in A and B. (D) Boxplot highlighting the difference between the percentage of genes showing inversion for GSTF pairs within the orange branch (Fig 2A), GSTF pairs outside this cluster and K/P pairs. p values are based on a two-sided Mann-Whitney test.

### Inversion is associated with a specific subset of GSTFs

Within the slow growth corrected genetic interaction profiles another interesting cluster stands out: the orange branch where five out of six pairs involve GSTFs which predominantly show the inversion pattern (Fig 2A, branch 3). This suggests that inversion may be strongly associated with a particular group of GSTFs, whereas this does not seem to be the case for kinases and phosphatases. The overall percentage of genes showing inversion is already much higher for GSTFs (28.6%) than for kinases/phosphatases (18.7%) (S2 Table). When investigating the GSTF pairs within the cluster, it is clear that these display an even higher percentage of inversion compared to kinases and phosphatases (Fig 2D; p=0.00026) as well as compared to other GSTF pairs (Fig 2D; p= 0.0043). In order to determine whether inversion was specific to the set of GSTFs analyzed here, or part of a more general phenomenon common to GSTFs, we included both positive and negative genetic interactions between GSTF pairs, expanding the number of GSTF pairs to 44. Clustering of all 44 GSTF pairs (S4 Fig) also shows that a large fraction of the GSTF pairs contain many genes showing inversion, with most of the inversion dominated GSTF pairs still clustering together (S4 Fig, indicated with an asterisk). Note though, that because the 44 GSTF pairs include both positive and negative genetic interactions, the results are not directly comparable to the kinase/phosphatase pairs as these only include negative genetic interactions. Taken together, this indicates that not only is inversion more frequently associated with GSTFs compared to kinases and phosphatases, but one particular subset of GSTFs is also predominantly defined by inversion.

### An exhaustive modeling approach to explore potential mechanisms underlying inversion

Unlike buffering, where redundancy is a likely mechanistic explanation, the underlying mechanism of inversion is still unknown [17]. The GSTF pairs within the inversion dominated cluster also do not share a common biological process, function, pathway or protein domain other than general transcription related processes and functions. To investigate potential mechanisms of inversion, an exhaustive exploration was initiated. Previously, Boolean modeling has been applied to exhaustively explore all mechanisms underlying two genetic interaction patterns for the Fus3-Kss1 kinase phosphatase pair [16]. However, to explore all potential mechanisms underlying inversion, a Boolean approach may not suffice as more subtle, quantitative effects, may be needed to obtain inversion. At the same time, any modeling approach must remain computationally feasible. For this purpose, a modeling approach based on Petri nets was devised to exhaustively evaluate all possible three and four node models but taking into account the possibility of quantitatively different effects (Fig 3, Methods). Interactions between nodes (edges) can be activating (positive) or inhibiting (negative). In order to incorporate quantitative differences, both strong and weak edges were used (Methods). Counting all possible combinations of different edges results in 152,587,890,625 possible edge weight matrices. To reduce the number of models, three conditions were imposed, as used previously [16]. In short, nodes contain no self-edges, the number of incoming edges on any node is limited to two and the model includes at least two edges from one of the regulators (R1, R2) to the downstream genes (G1, G2). Applying these requirements and filtering for mirror edge weight matrices results in 2,323,936 matrices. By including AND/OR logics the final number of models to be evaluated was 9,172,034 (Methods). Petri net simulations were then run and genetic interaction patterns determined for G1 and G2, analogous to what was done for the original data (Methods) (Fig 1A). Depending on the topology, Petri net models can be stochastic, in other words, they do not show the same behavior when simulated multiple times and therefore result in unstable models. Only 2.3% of the models were found to be unstable, i.e. showed inconsistent genetic interaction patterns for G1 and G2 across five times simulation runs. Thus, stochasticity hardly influences the observation of genetic interaction patterns in our simulations (Fig 3). Nevertheless, unstable models were excluded from further analysis. In total, 168,987 models (1.8%) show inversion in either G1, G2, or both downstream nodes.

**Fig 3.**
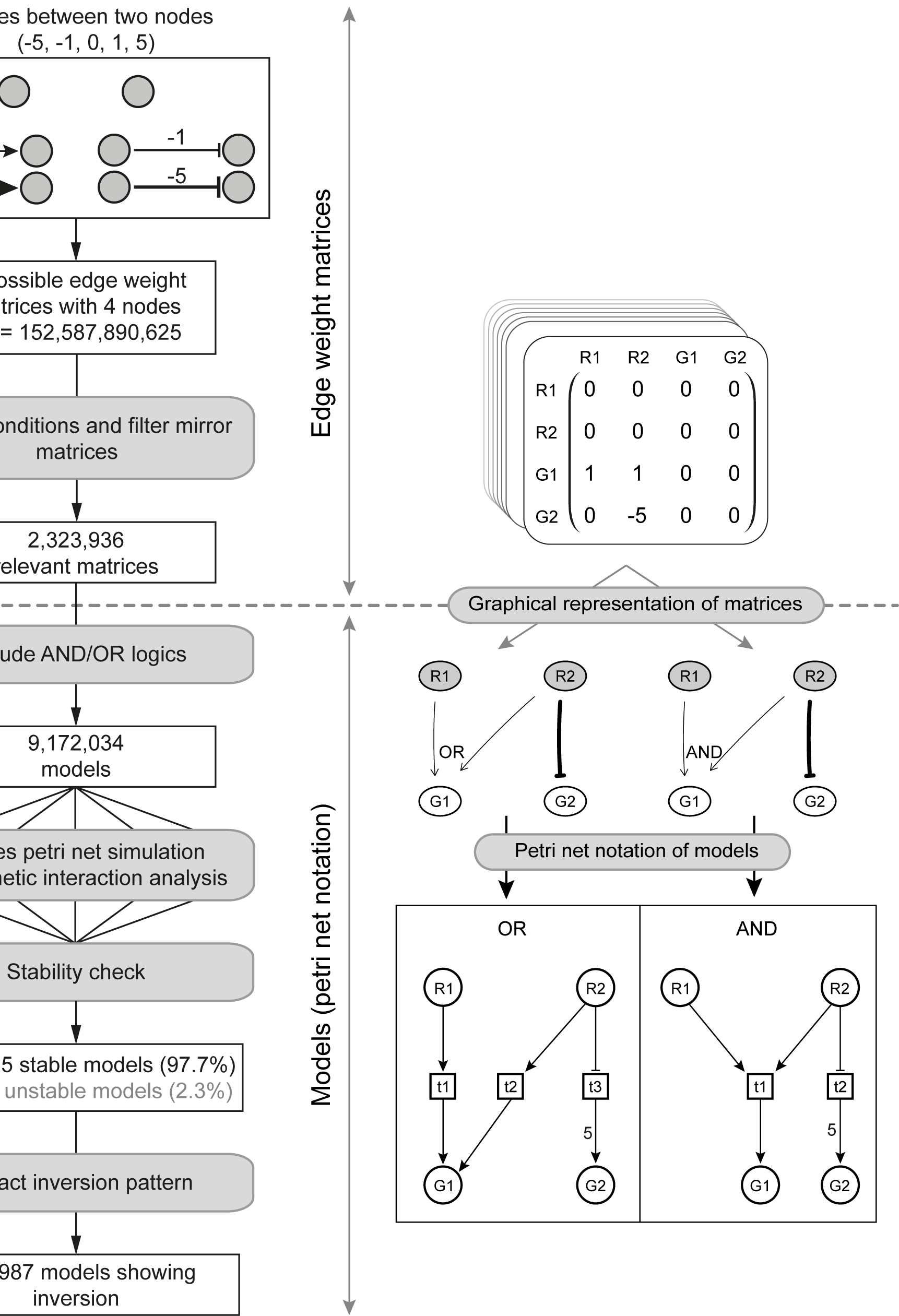
Schematic overview of Petri net simulation pipeline

Schematic overview of the pipeline implemented for performing Petri net simulations. The left panels show from top to bottom the different steps performed when running the simulation pipeline. The right panels show the different data representations used throughout the pipeline. The right panel above the dashed line indicates a series of steps where edge weight matrices are used. The right panel below the dashed line indicates steps where models or Petri net notation are used.

### A quantitative difference in interaction strength is a strict requirement when observing inversion

To investigate which potential regulatory patterns underlie the 168,987 models showing inversion, low complexity models with few edges were analyzed first. Two interesting observations can be made. First, although there are many high complexity models involving four nodes and many edges (up to eight), three nodes and three edges are sufficient to explain inversion (Fig 4A). Second, only two three-node models exist that show inversion (Fig 4A). These two models only differ in the strength of the inhibiting edge from R1 to R2. Both models involve inhibition of R2 through R1 and weak activation of G1 by R1 in combination with a strong activation of G1 by R2, i.e. a quantitative edge difference between the incoming edges of G1. Deletion of R1 in these two models results in activation of R2, and therefore upregulation of G1 due to a strong activating edge. Deletion of R2 however, will not result in any changes compared to WT as it is normally inhibited by R1. Deletion of both R1 and R2 will lead to downregulation of G1 as the weak activating edge from R1 to G1 is lost. Taken together, the analysis of the low complexity models indicates that a quantitative difference in interaction strength is required to explain inversion. To investigate whether this requirement also holds for higher complexity models, all models containing two to eight edges were further analyzed. Inversion models were grouped by the number of edges (complexity) and then analyzed for their relative frequency of having a quantitative edge difference (Fig 4B, top left panel, note that the number of possible models grows exponentially with the number of edges). Almost all of these models show a quantitative edge difference, with only a very small fraction (1.3% overall) of models not having a quantitative edge difference. Except for masking, the other genetic interaction patterns show different behavior, indicating that the relative ratio of quantitative versus non-quantitative edges is not an inherent network property. Based on both the low complexity models as well as the high complexity models showing inversion, it is evident that a quantitative difference in interaction strength of two genes or pathways acting on a downstream gene is required to explain inversion.

**Fig 4.**
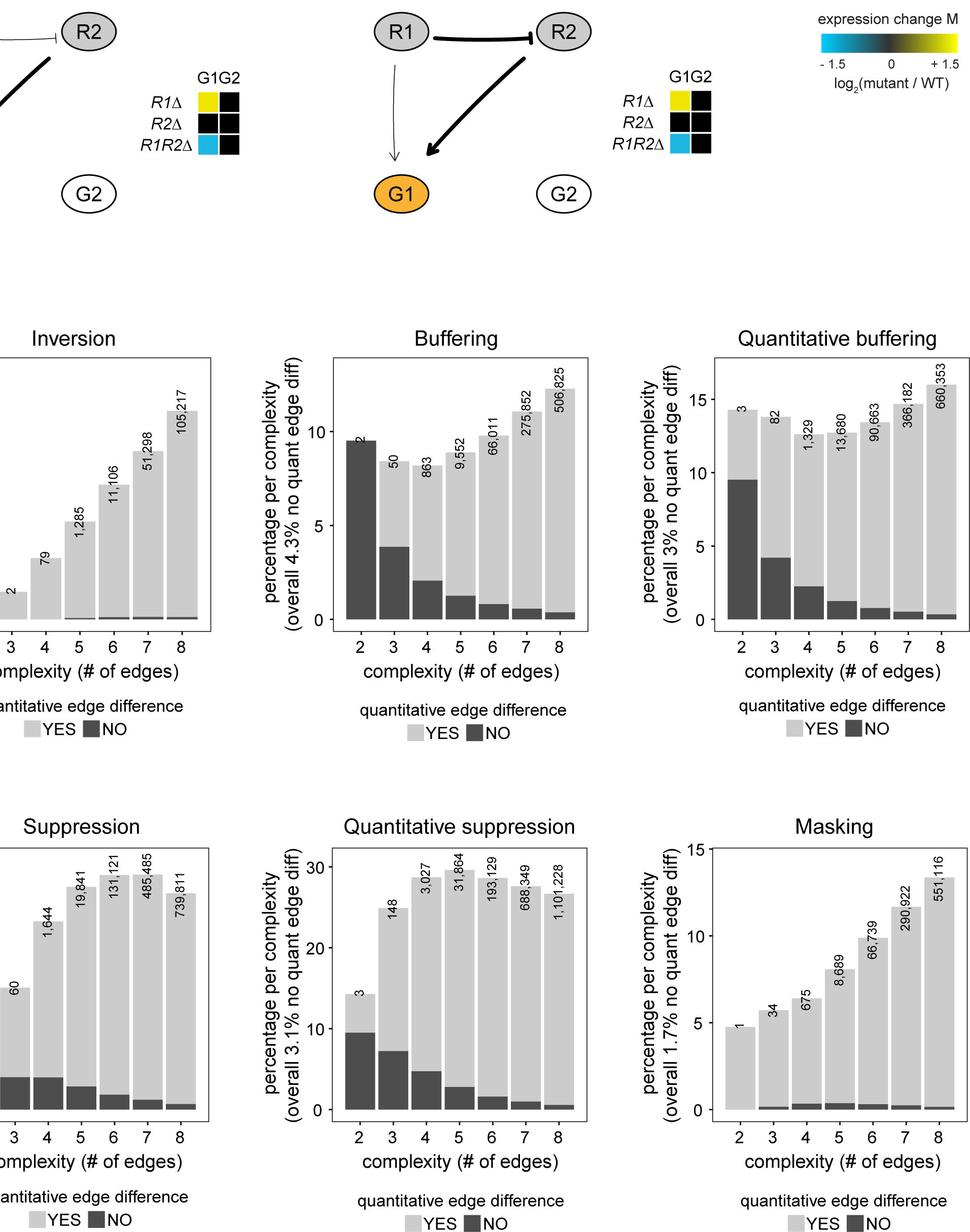
A quantitative edge difference is the minimum requirement for observing inversion. (A) Petri net simulation results for the only two models with three nodes that result in inversion (indicated in orange) for the G1 node. Heat maps indicate the log_2_(FC) of the number of tokens in simulated deletion mutants (single and double mutant) relative to the WT situation. Thicker lines indicate edges with a strong effect. (B) For each genetic interaction pattern (inversion, buffering, quantitative buffering, suppression, quantitative suppression and masking), the percentage of models showing that particular genetic interaction pattern is shown, split up per complexity (number of edges). The percentage per complexity is calculated as the number of models showing a particular genetic interaction pattern for a certain complexity, divided by the total number of models for that complexity. Bar plots are subdivided into two types of models, models that have quantitative differences between edge weights (bright gray) and models that have no quantitative differences between edge weights (dark gray). The number of models showing the particular genetic interaction pattern per complexity is shown on top of each bar plot.

### A quantitative difference in interaction strength is frequently accompanied by an intermediate buffering node

With the exception of the two models discussed above, all other inversion models consist of four nodes with two regulator nodes and two downstream effector nodes. To better understand the interplay between all four nodes, besides the node displaying inversion (G1), the second downstream gene (G2) was also analyzed for the occurrence of different genetic interaction patterns (Fig 5A). Most G2 nodes tend to have no genetic interaction pattern (27%). The most common genetic interaction patterns are buffering (23%) and quantitative buffering (18%). These both are very alike in their genetic interaction pattern (Fig 1A) and only show slight differences in their quantitative behavior. They may therefore be considered as part of the same superclass of “buffering”. The buffering node is frequently positioned upstream of the inversion node, and always downstream of R1/R2 (Fig 5B). The combination of inversion and buffering is also significantly overrepresented within inversion models when compared to all models (Table 1, p < 0.005). Taken together this shows that a quantitative difference in interaction strength of two genes or pathways acting on a downstream gene is frequently accompanied by an intermediate gene or pathway that displays buffering.

**Fig 5.**
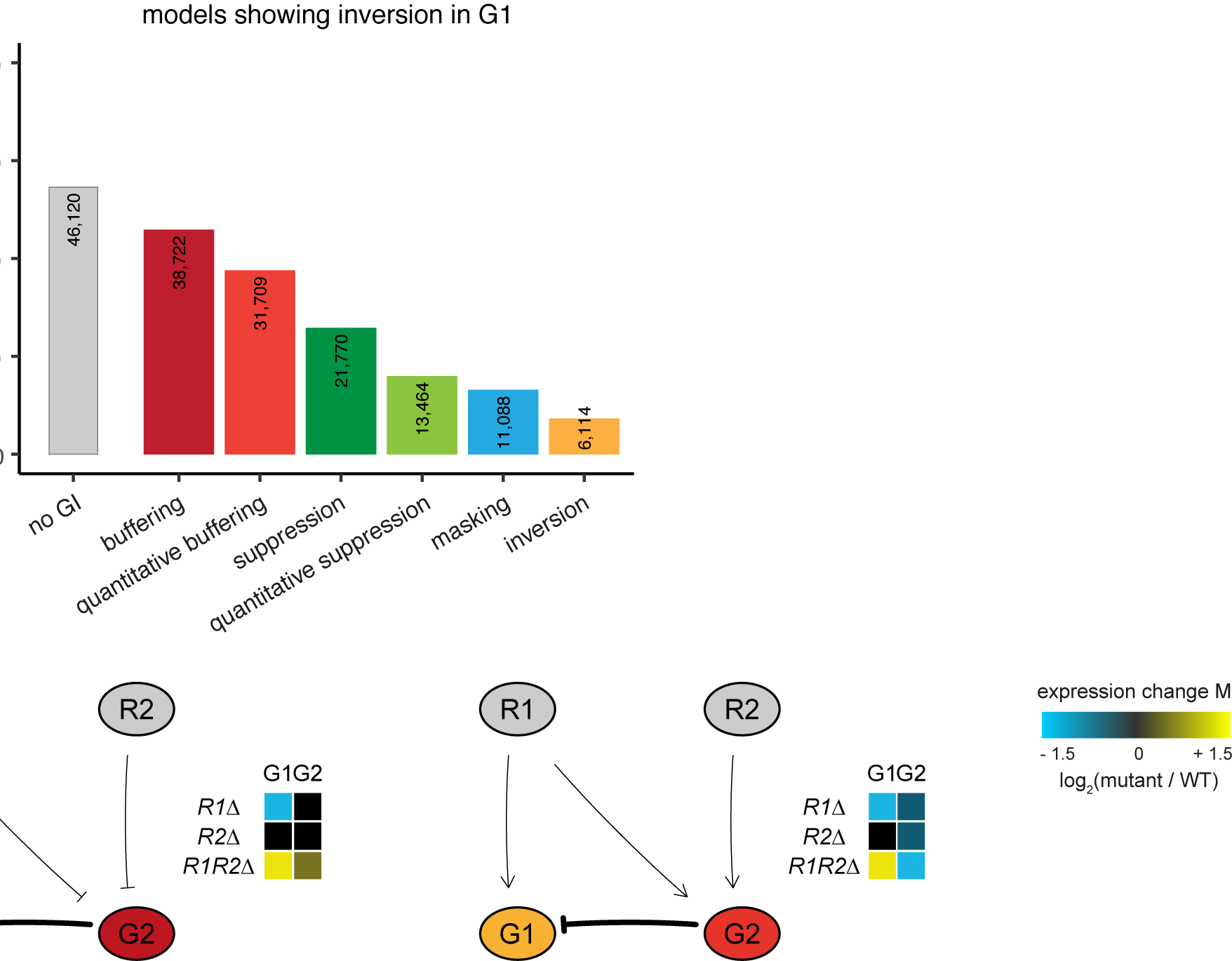
Inversion is frequently accompanied by buffering. (A) Bar plots showing the percentage of models that either have no genetic interaction (gray, left bar) or a different genetic interaction pattern in node G2 when node G1 is displaying inversion. The number of models per category is shown on top of each bar plot. Color scheme of the genetic interaction patterns as in Fig 1A. (B) Petri net simulation results for two models with four nodes with node G1 always displaying inversion and node G2 displaying either buffering (left) or quantitative buffering (right). Heat maps as in Fig 4A.

**Table 1.**
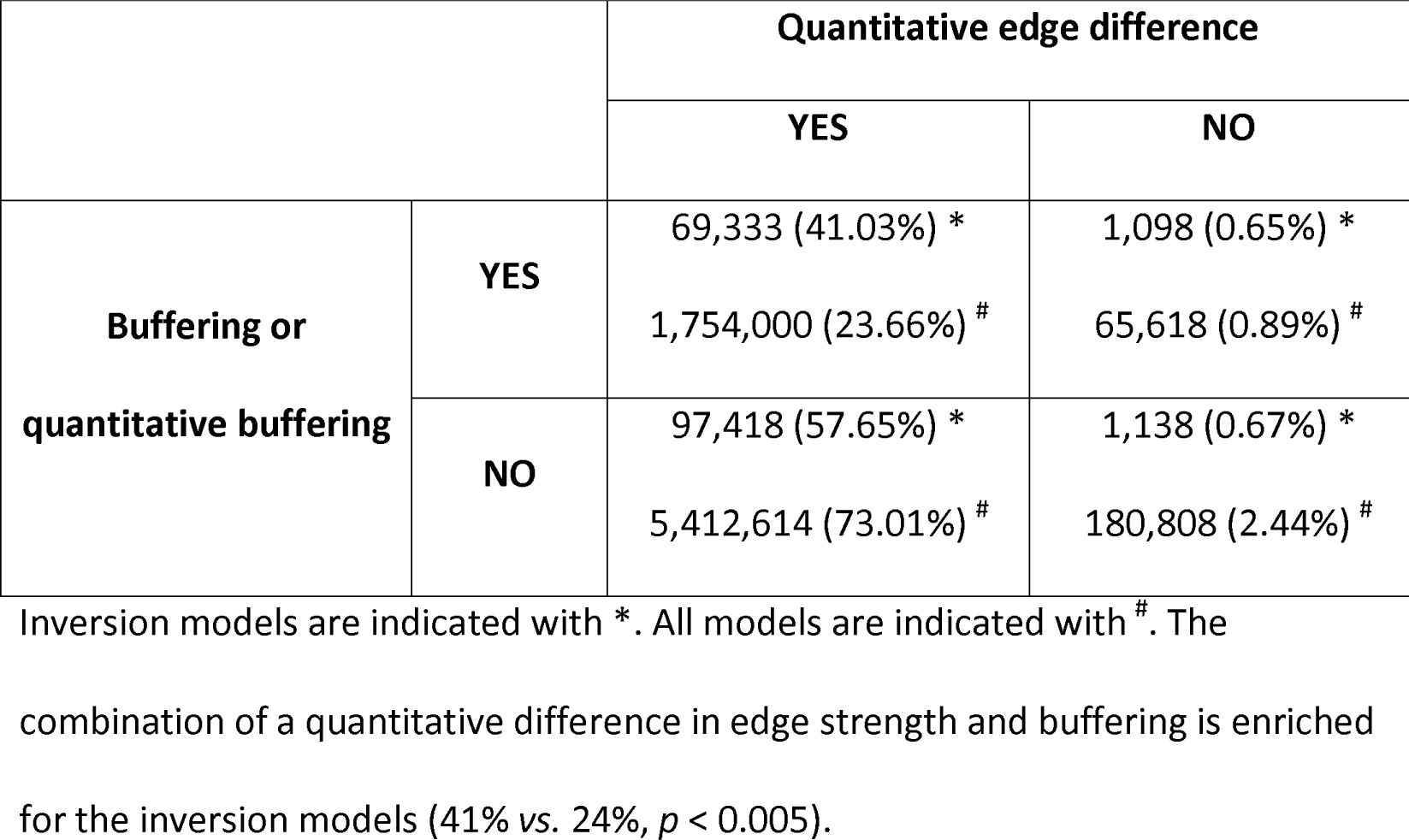
Models with a quantitative edge difference and intermediate buffering node

### Gat1 and Gln3 might differentially regulate mitochondrial-to-nuclear signaling

One gene pair within the inversion dominated GSTF cluster (Fig 2A, branch 3; Fig 6A) that largely consists of inversion is Gat1-Gln3. By combining the three node model derived from the Petri Net modelling (Fig 4A, left panel) with existing literature, a potential mechanistic explanation for the interaction between this pair can be obtained (Fig 6B). Both Gln3 and Gat1 are activators involved in regulating nitrogen catabolite repression (NCR) sensitive genes [62–64]. When cells are grown under nitrogen rich conditions, as was done here, Gat1 is repressed by Dal80 [63]. Dal80 in turn can be activated by Gln3 [63,65], which provides a plausible mechanism for the predicted inhibition edge between Gln3 and Gat1 (Fig 6B). The degree to which Gln3 and Gat1 influence downstream genes has also been reported to differentiate between individual genes [66], which is fully consistent with the quantitative edge difference as predicted in the model (Fig 6B). The set of inversion related genes (Fig 6A, gene set 1) is enriched for nuclear encoded mitochondrial respiratory genes (Fig 6A, denoted with a dot, p value 3.2×10^-17^). Previously, NCR has been linked with mitochondrial-to-nuclear signaling through the retrograde signaling pathway [67,68], although an alternative mitochondrial-to-nuclear signaling pathway, such as the intergenomic signaling pathway, may instead be involved [69]. Taken together, this suggests that Gat1 and Gln3 might differentially influence mitochondrial-to-nuclear signaling, although additional experiments would be needed to confirm this initial hypothesis.

**Fig 6.**
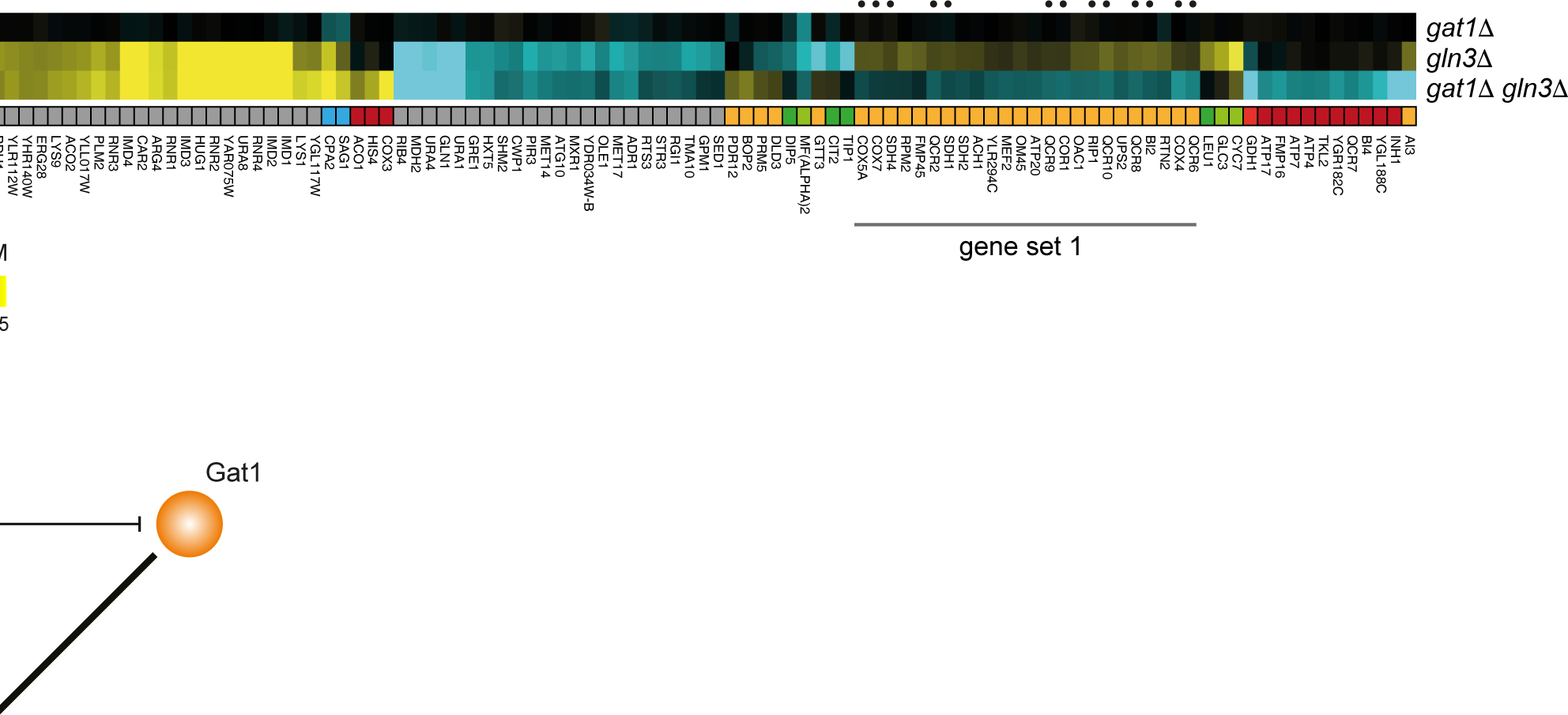
Gln3 and Gat1 might differentially regulate mitochondrial-to-nuclear signaling. (A) Expression changes compared to WT (horizontal) in *gat1Δ, gln3Δ, and gat1Δ gln3Δ* mutants (vertical) after slow growth correction. Different colors underneath the gene expression profiles represent different genetic interaction patterns as indicated in Fig 1A. Gray depicts gene expression changes not part of a genetic interaction pattern. Nuclear encoded mitochondrial respiratory genes are denoted with a dot. (B) Proposed model to explain the inversion pattern between Gat1 and Gln3 based on the Petri net simulation result in Fig 4A.

### Pdr3 likely acts as the intermediate buffering gene in mediating the inversion pattern observed for Hac1-Rpn4

Another interesting pair of genes within the GSTF cluster dominated by the inversion pattern (Fig 2A, branch 3) is Hac1-Rpn4. This pair displays a substantial amount of both inversion as well as buffering (Fig 7A) and lends itself well for testing some of the model predictions. Hac1 and Rpn4 are both involved in the processing of inappropriately folding proteins, either by activating genes of the unfolded protein response [70] (UPR, Hac1) or via the endoplasmic reticulum-associated degradation [71] (ERAD, Rpn4). Two genes that display inversion, Pdr5 and Pdr15, show stronger expression changes compared to the other genes in the same gene set (Fig 7A, gene set 1). Both Pdr5 and Pdr15 are multidrug transporters involved in the pleiotropic drug response [72]. Expression of these two genes is tightly regulated by Pdr1 and Pdr3 [73,74]. Pdr5 is also positively regulated by expression of Yap1, a basic leucine zipper transcription factor that is required for oxidative stress tolerance [75]. Of the three transcription factors Pdr1, Pdr3 and Yap1, only PDR3 shows a clear upregulation in the *hac1Δ rpn4Δ* double mutant and hardly any change in the respective single mutants (Fig 7B). This is consistent with the role of the intermediate buffering gene as derived from our Petri net modelling results. If Pdr3 acts as the intermediate buffering gene as predicted based on our model, it is also expected that deletion of PDR3 leads to a more severe downregulation of PDR5 and PDR15 expression levels when compared to expression levels of PDR5 and *PDR15* in the *rpn4Δ* mutant. To test this prediction, mRNA expression changes of PDR5 and PDR15 where investigated in the *pdr3Δ* and *rpn4Δ* mutants. As expected, deletion of PDR3 results in a much stronger downregulation of PDR5 (p=7.26×10^-4^) and PDR15 (p=5.95×10^-5^) compared to deletion of RPN4 (Fig 7C), thereby confirming the model prediction. Taken together, these results provide a likely mechanistic explanation where Pdr3 acts as the intermediate buffering gene in regulating Pdr5 and Pdr15 (Fig 7D).

**Fig 7.**
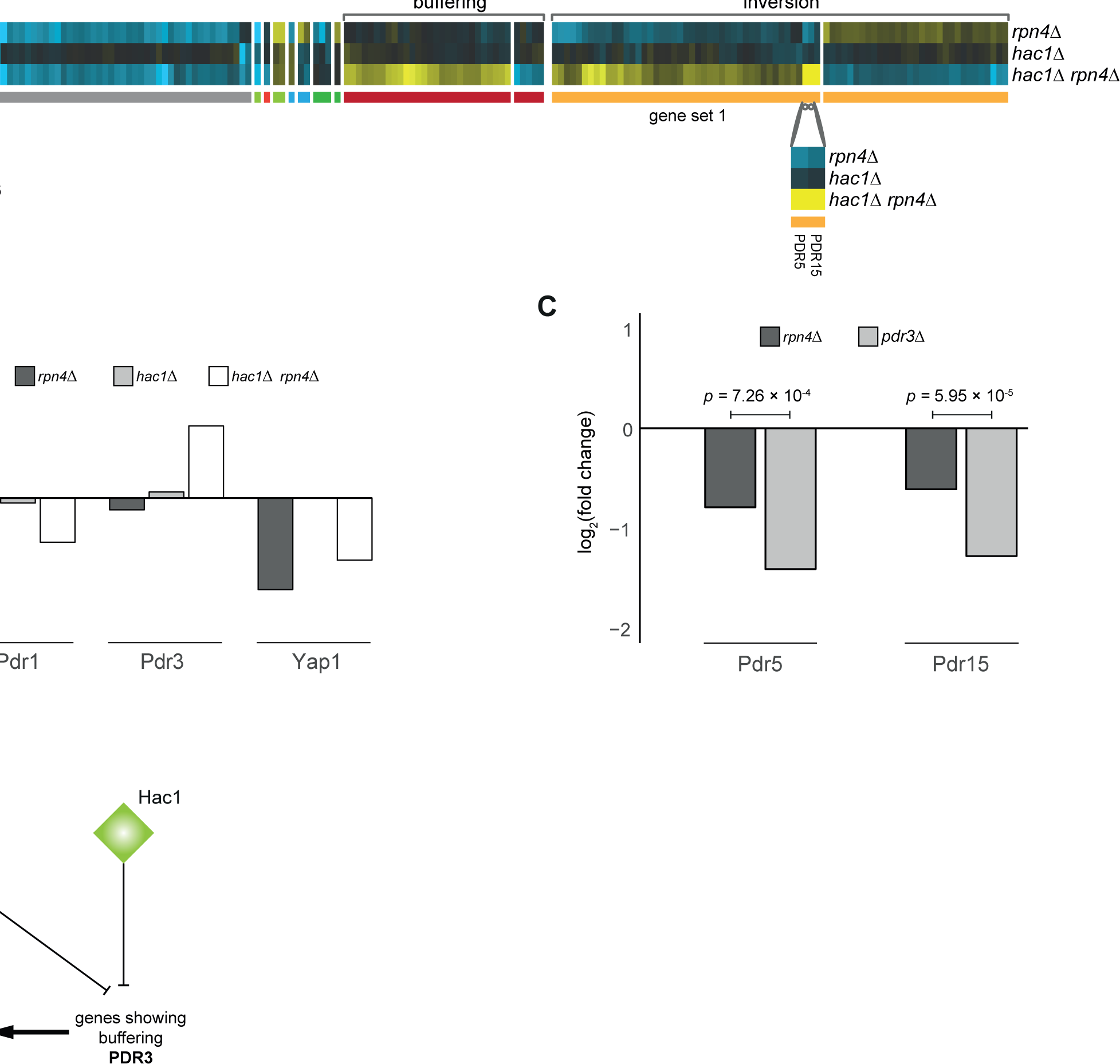
Pdr3 acts as an intermediate gene for observing inversion in PDR5 and PDR15. (A) Expression changes compared to WT (horizontal) in rpn4Δ, hac1Δ, and hac1Δ rpn4Δ mutants (vertical) after slow growth correction. Different colors underneath the gene expression profiles represent different genetic interaction patterns as indicated in Fig 1A. Gray depicts gene expression changes not part of a genetic interaction pattern. (B) Expression changes of Pdr1, Pdr3 and Yap1 compared to WT in rpn4Δ, hac1Δ and hac1Δ rpn4Δ mutants. (C) Expression changes of Pdr5 and Pdr15 compared to WT in rpn4Δ and pdr3Δ mutants. P values are obtained from a limma analysis comparing gene expression changes between rpn4Δ and pdr3Δ mutants. (D) Proposed model to explain the inversion pattern between Hac1 and Rpn4 based on the Petri net simulation result in Fig 5B.

## Discussion

### Genome-wide gene expression measurements to investigate the genetic interaction landscape

To investigate genetic interactions in a high-throughput manner, growth-based assays have frequently been deployed, resulting in the identification of an overwhelming number of both negative and positive genetic interactions [6,20–28]. Based on these surveys, several theoretical mechanisms have been proposed to explain genetic interactions [3,18,76,77]. More efforts, also using different types of assays, are however still needed to systematically and thoroughly investigate the underlying mechanisms. Alongside growth-based genetic interactions, genome-wide gene expression measurements have been applied to elucidate potential molecular mechanisms underlying genetic interactions [16,17,33–36]. Although more laborious, expression-based genetic interactions potentially allow for more in-depth characterization of the genetic interaction landscape. Here, we show that buffering is the most frequently occurring pattern underlying most negative genetic interactions. These are however to a large degree related to slow growing strains, hindering the investigation of the underlying mechanisms. By applying a slow growth transformation that removes a cell cycle associated gene expression signature, many such effects can be filtered out [48]. The transformation results in distinct clusters that can be more easily aligned with potential underlying mechanisms. Recent advances using Crispr-Cas9 single and double knock-down screens, followed by single cell RNA sequencing have also shown that results are greatly influenced by the cell-cycle phase in which different cells are found [35,78]. It is therefore essential for future studies on genetic interactions to incorporate methods that decompose such large confounding effects, as they greatly influence the ability to deduce mechanism.

### Systematic modelling to understand mechanisms of genetic interactions

To infer underlying mechanisms from the genetic interaction landscape as obtained from genome-wide gene expression measurements, systematic modeling approaches are warranted [3,18]. Various modeling techniques have been instrumental in understanding various aspects of experimental data (reviewed in [79]). Different modeling methods have different applications, depending on the question asked and available data types. To infer the underlying mechanisms for many genetic interactions, an approach is needed that is able to exhaustively explore the complete genetic interaction landscape while at the same time incorporating (semi-) quantitative values. Here, using Petri net modeling, we have been able to exhaustively explore more than nine million models that included semi-quantitative effects. Inversion, a pattern strongly associated with a group of GSTF pairs was investigated in more detail, resulting in the striking conclusion that a quantitative difference in interaction strength is needed to explain inversion. The approach taken here, by combining slow growth corrected genome-wide gene expression measurements with the exhaustive semi-quantitative Petri-net modeling thus highlights the benefits of using such an approach to understand mechanisms of genetic interactions. Applying this approach to other types of genetic interactions or across many more genetic interaction pairs can help us in further characterizing mechanisms of genetic interactions and relating these to pathway organization and cellular states.

### Inversion as a way to differentially regulate between two redundant processes and a third, compensatory process

Previously, a mechanism termed “buffering by induced dependency” was proposed to explain parts of the genetic interaction patterns observed between Rpn4 and Hac1 (Fig 8, dotted inset) [17]. This mechanism links the endoplasmic reticulum-associated degradation (ERAD) by the proteasome (Rpn4) with the unfolded protein response (UPR, Hac1), two distinct processes dealing with misfolded and unfolded proteins. By combining the “buffering by induced dependency” mechanism with the model proposed for inversion here, most genetic interaction patterns observed for Rpn4 and Hac1 can be explained (Fig 7A; 8). The combined model introduces a third, compensatory process, the pleiotropic drug response (PDR; Fig 8, bottom light gray inset). Even though the exact relationship between ERAD, UPR and pleiotropic drug response is unclear, the interplay between UPR and drug export has been shown in mammalian cells [80]. In yeast, Pdr5 and Pdr15 have been implicated in cellular detoxification [74,81] and may also be required for cellular detoxification under normal growth conditions [81]. Both Pdr5 and Pdr15 have been reported to be regulated through Pdr1 and Yap1 [75,82], as well as through Rpn4 [83,84]. This is also confirmed here by downregulation of both Pdr1 and Yap1 as well as downregulation of their target genes Pdr5 and Pdr15 in *rpn4Δ* (Fig 7B, C). It is therefore likely that in the wildtype situation when Rpn4 is active, both ERAD and the PDR are functioning (Fig 8). Deletion of RPN4 leads to deactivation of the ERAD and PDR pathways and activation of the UPR through Hac1 (Fig 8, *rpn4Δ* dotted red line). Deletion of both RPN4 and HAC1 results in a major growth defect and accumulation of misfolded and unfolded proteins, most likely leading to a stronger activation of the PDR through Pdr3 compared to the wildtype situation (Fig 7B, C; Fig 8, *hac1Δ rpn4Δ* dotted red line) [73,74]. Taken together, this model thus provides a potential regulatory mechanism in which two redundant processes, each with slightly different efficacies, can be differentially regulated, or fine-tuned, through a third, compensatory process. The requirement to fine-tune slightly different efficacies of different cellular processes then also provides a potential explanation why inversion is observed more frequently for gene-specific transcription factors since these allow for more fine-grained control than protein kinases and phosphatases.

**Fig 8.**
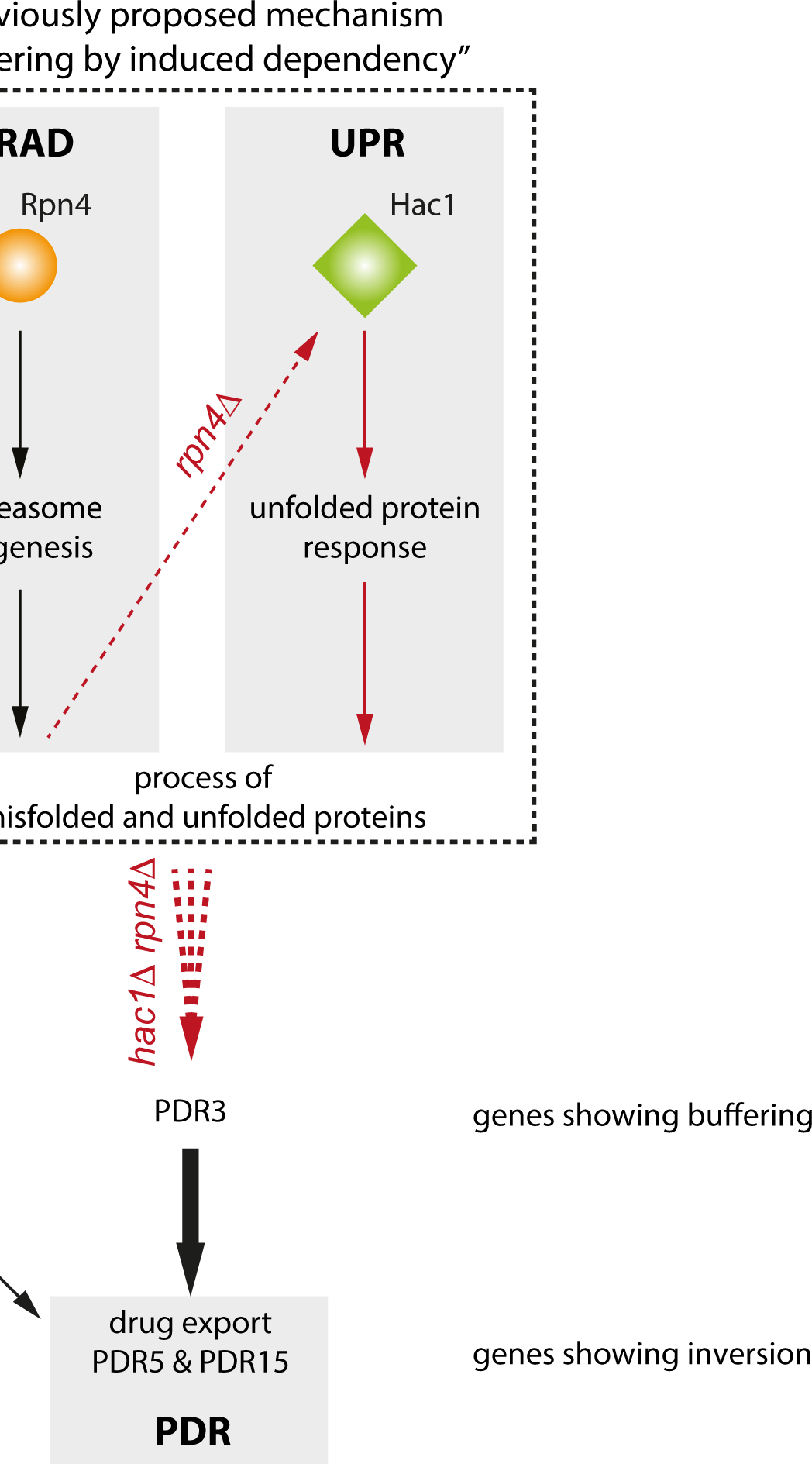
Combination of buffering by induced dependency and proposed model for inversion. Carton depiction of proposed model for genetic interaction between Rpn4 and Hac1. Red arrows indicate the consequence of disrupted genes and pathways. The dashed rectangle indicates a previously proposed model, “buffering by induced dependency”, to explain genes showing buffering for Hac1-Rpn4. A thicker arrow represents a stronger activation strength.

In conclusion, we have shown how exhaustive exploration of regulatory networks can be used to generate plausible hypothetical regulatory mechanisms underlying inversion. Almost all models showing inversion contain a quantitative difference in edge strengths, which suggests quantitative differences in regulating downstream target genes are important for the functioning of GSTFs. These hypothetical mechanisms have subsequently been tested against known and new experimental data. For GSTFs we show a validated example of Hac1-Rpn4 where differential regulation of gene expression is key to understanding the genetic interaction pattern inversion.

## Materials and Methods

### Selection of GSTF and kinase/phosphatase pairs

Two selection criteria were applied to select genetically interacting GSTF and kinase/phosphatase pairs. First, one of the mutants of each individual pair should show genome-wide gene expression measurements similar to wildtype (WT). DNA microarray data from Kemmeren et al [85] was used to determine whether a single deletion mutant is similar to WT. A deletion mutant is considered similar to WT when fewer than eight genes are changing significantly (p < 0.05, FC > 1.7) in the deletion mutant gene expression profile, as previously described [16]. Second, selected pairs should show a significant growth-based negative genetic interaction score. Growth-based genetic interaction scores for GSTF [28] and kinase/phosphate [26] pairs were converted to Z-scores. A negative Z-score significance of p < 0.05 after multiple testing correction was used as the significance threshold. Applying these selection criteria resulted in 11 GSTF pairs and 15 kinase/phosphatase pairs (S1 Table).

### Genome-wide gene expression measurements and statistical analyses

Genome-wide gene expression measurements of single and double mutant GSTF pairs were obtained from Sameith et al [17]. Genome-wide gene expression measurements of single and double mutant kinase/ phosphatase pairs were obtained from van Wageningen et al [16]. Genome-wide gene expression measurements of *pdr3Δ* and *rpn4Δ* were obtained from Kemmeren et al [85]. Statistical analysis of these gene expression profiles was performed as previously described [85]. In summary, mutants were grown in Synthetic Complete (SC) medium with 2% glucose and harvested during exponential growth. WT cultures were grown alongside mutants in parallel to monitor for day to day effects. For each mutant statistical analysis using limma was performed versus a collection of WTs [16,85]. Reported FC for each transcript is the average of four replicate expression profiles over a WT pools consisting of 200 WT strains.

### Growth-based genetic interaction scores

Growth measurements for single and double mutant GSTF and kinase/phosphatase pairs were obtained from Sameith et al [17] and van Wageningen et al [16] respectively. Growth-based genetic interaction scores were calculated for both GSTF and kinase/phosphatase pairs as performed before [17]. In summary, the fitness W of single and double mutants was determined as the ratio between the WT growth rate and the mutant growth rate. The growth-based genetic interaction score 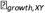 was calculated as the deviation of the observed fitness in a double mutant from the expected fitness based on the respective single mutants 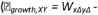 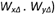. *P* values were assigned to genetic interaction scores based on the mean and standard deviation of a generated background distribution [17]. *P* values were corrected for multiple testing using Benjamini-Hochberg. Adjusted *p* values lower than 0.05 were considered significant. Fitness values of all single and double mutants, as well as calculated genetic interaction scores can be found in S1 Table.

### Expression-based genetic interaction scores

Expression-based genetic interaction scores were calculated for both GSTF and kinase/phosphatase pairs as described before [17]. In summary, the effect of a genetic interaction between two genes X and Y on gene i is calculated as the deviation between the observed expression change in the double mutant and the expected expression change based on the corresponding single mutants (ε*_txpn_i,XY_* = |M_*i,xΔyΔ*_ - (M_*i,xΔ*_ + M*_i,yΔ_*) |). The overall genetic interaction score between X and Y is calculated as the sum all genes i for which *ε_txpn_i,XY_* > log2(1.5). All genetic interaction scores consisting of at least 10 genes were kept for further downstream analyses. Genes with similar gene expression changes were divided into the 6 different patterns (buffering, quantitative buffering, suppression, quantitative suppression, masking, inversion), as previously described [17] (Fig 1A).

### Clustering of expression-based genetic interaction scores

Genetic interaction profiles for both classes of proteins were grouped together based on the number of occurrences of the six different patterns using hierarchical clustering. Average linkage was applied for the clustering. Identity of genes in each genetic interaction profile was disregarded.

### Slow growth transformation

Slow growth signature transformation of the gene expression profiles was performed as previously described [48]. In short, for each mutant, the correlation of its expression profile with the first principal component of 1,484 deletion strains [85] was removed, thus minimizing correlation with the relative growth rate. The transformation reduces correlation with the relative growth rate from 0.29 to 0.10 on average [48].

### Model generation

Exhaustive modeling of possible network topologies underlying the genetic interaction patterns was carried out by creating Petri net models consisting of four nodes, representing two regulator genes (R1 and R2) and two downstream genes (G1 and G2). With four nodes and directed edges, there are 4^2^=16 possible edges, and 2^16^=65536 possible edge weight matrices, which is a tractable number. However, each interaction can in addition be positive or negative, and weak or strong (and absent), leading to 5^16^=1.5010^11^ possible interaction graphs (edge weight matrices), which becomes intractable. Many of these models, however, will be irrelevant for the understanding the biological behavior of genetic interaction patterns of two genes. To exclude these types of models, the following conditions were applied: 1) No self-edges are allowed. 2) The number of incoming edges on any node must be limited to two. 3) At least two incoming edges from at least one of the regulators (upstream nodes) to the genes (downstream nodes). Applying these conditions reduces the number of relevant edge weight matrices to 9,287,616. Furthermore, most generated matrices have mirror counterparts, therefore only one of the matrices was included in downstream analyses. Applying this filtering step results in 2,323,936 matrices. Fig 3 gives an overview of the various filtering steps, and shows which representation of the models was relevant in different stages of the filtering. Edge weight matrices were generated in R, version 3.2.2 (the function expand.grid was used to generate all combinations of edges per row in a given matrix).

### Petri net simulations

Regulatory effects of two potentially interacting genes (R1 and R2) on two downstream genes (G1 and G1) were simulated using a Petri net approach [42,44,86,87] to recapitulate genetic interaction patterns observed in the gene expression data.

In the Petri net notation, nodes in a given model are represented by places (denoted as circles). Interactions between nodes always go via a transition (denoted as squares), connected via directed arcs (drawn as arrows). An incoming arc to a transition can be either activating or inhibiting. The weight on arcs going to a transition is always fixed to 1. The weight on arcs going from a transition to a place depends on the edge weight between two nodes, 1 for weak and 5 for strong (Fig 3).

For nodes with two incoming edges, one has to decide how these two inputs should be combined: does the transition require both inputs to be activated (AND logic), or can one or the other activate it (OR logic). To incorporate this, for each pair of incoming edges with the same weight, two Petri net models were generated: one using the AND logic, and one using the OR logic (Fig 3, bottom right panel). For two incoming edges with different weights only the Petri net model using the OR logic was generated. For cases with two incoming edges to a node with two different directions, activation and inhibition, inhibition dominates.

To simulate the regulatory effects of two upstream genes (R1 and R2), 200 tokens were provided to represent the mRNA resources for each regulator, except when one of the regulators has an incoming edge from the other regulator as shown in (S5A Fig). Each step in the simulation process comprises of firing all enabled transitions (maximal parallel execution) [88,89]. A transition is enabled to fire when resources (tokens) in the input place(s) match or exceed the weight(s) on the respective incoming arc(s) to the transition (S5B Fig). In total 50 consecutive transition firing steps were performed.

To incorporate deletion mutants in the simulation process, tokens were removed from corresponding regulators. To prevent accumulation of tokens in deleted regulators, each outgoing arc from a transition to the corresponding deleted places were also removed in simulated deletion strains. The number of tokens in G1 and G2 after 50 steps of firing transitions in single and double mutants were compared with that in the WT situation where both R1 and R2 are active. To avoid division by zero one token was added to the total number of tokens in G1 and G1. These fold changes were then log2 transformed (M values).

Simulation-based genetic interaction scores for G1 and G2 were calculated based on the deviation between observed M values in the double mutant and the expected M value based on the single mutants, as follows: ε_*sim,R1R2i*_ = |M_R1ΔR2Δi_ - (M_*R1Δi*_ + M_*R2Δi*_)|, where *i* can be either G1 or G2. Each node with ε_*sim,R1R2i*_ > log2(1.7) was further divided into genetic interaction patterns, as defined before based on gene expression data [17]. Simulated expression levels for single and double mutants are considered to be increased relative to WT when M > log2(1.7) and decreased when M < -log_2_(1.7).

### Functional enrichment tests

Functional enrichment analyses were performed using a hypergeometric testing procedure on Gene Ontology (GO) biological process (BP) annotations [67] obtained from the *Saccharomyces Cerevisiae* Database [68]. The background population of genes was set to 6,359 and p values were corrected for multiple testing using Bonferroni.

### Visualization of models

Models were visualized in R, version 3.2.2, using diagram package (version 1.6.3). Weak and strong activation/inhibition edges are represented as thin and thick lines, respectively.

## Acknowledgements

We would like to thank Wim de Jonge, Thanasis Margaritis and all the other lab members for helpful discussions and advices throughout the project.

## Author Contributions

**Conceptualization**: Saman Amini, Frank C. P. Holstege, K. Anton Feenstra, Patrick Kemmeren

**Formal Analysis**: Saman Amini, Annika Jacobsen, Olga Ivanova, Philip Lijnzaad

**Funding Acquisition**: Patrick Kemmeren

**Investigation**: Saman Amini, Annika Jacobsen, Olga Ivanova, Philip Lijnzaad, Jaap Heringa, Frank C. P. Holstege, K. Anton Feenstra, Patrick Kemmeren

**Methodology**: Saman Amini, Annika Jacobsen, Olga Ivanova, Frank C. P. Holstege, K. Anton Feenstra, Patrick Kemmeren

**Resources**: Frank C. P. Holstege, Patrick Kemmeren

**Software**: Saman Amini, Annika Jacobsen, Olga Ivanova

**Supervision**: Jaap Heringa, Frank C. P. Holstege, K. Anton Feenstra, Patrick Kemmeren

**Visualization**: Saman Amini, Annika Jacobsen, Olga Ivanova

**Writing** – Original Draft preparation: Saman Amini, Patrick Kemmeren

**Writing** – Review & Editing: Annika Jacobsen, Olga Ivanova, Philip Lijnzaad, Jaap Heringa, Frank C. P. Holstege, K. Anton Feenstra, Patrick Kemmeren

## Supporting Information

**S1 Table. Single and double mutant GSTF and kinase/phosphatase pairs**

**S2 Table. Number of genes for each genetic interaction pattern for both GSTF as well as kinases/phosphatase pairs.**

**S1 Fig. Buffering dominates genetic interaction profiles**

**(A)** Hierarchical clustering of all pairs according to their genetic interaction effects. Average linkage clustering was applied to group pairs with similar genetic interaction patterns. The number of occurrences for each genetic interaction pattern was used and the identity of individual genes was disregarded. Similarity between pairs was calculated using the cosine correlation. Most pairs are grouped together in a single branch (indicated in red), which is dominated by buffering. (B) The number of genetic interaction effects underlying the clustering are shown as bar plots below the dendrogram (top; colors as in Fig 1A). (B) Number of genes showing no genetic interaction pattern but significantly changing in one of the mutants compared to WT (bottom; p ≤ 0.01, FC > 1.5). Dark gray for the first named gene, light gray for the second named gene.

**S2 Fig. Slow growth correction improves identification of GSTF targets**

Scatter plots showing gene expression levels in the GSTF double mutant pairs hac1Δ rpn4Δ (A), met31Δ met32Δ (B), gat1Δ gln3Δ (C) and cbf1Δ hac1Δ (D) versus WT before (left) or after (right) slow growth correction. Individual transcripts are represented as dots. The dashed line indicates a FC of 1.7. Dots depicted in blue and red correspond to targets of the first and second gene in a named GSTF pair. P-values are calculated using a hypergeometric testing procedure to test the enrichment of GSTF targets among genes that change more than 1.7 fold before (left) or after (right) slow growth correction.

**S3 Fig. The genetic interaction between Elm1 and Mih1 can be explained through pathway redundancy**

Cartoon depicting the proposed genetic interaction between Elm1 and Mih1. (**left panel**) WT situation where the activity of Cdc28 is not disrupted by Swe1 phosphorylation. (**Middle panel**) Deletion of Elm1 leads to derepression of Swe1 activity. The increase of Swe1 activity can be compensated by Mih1. **(Right panel)** Deletion of both Elm1 and Mih1 will cause an increase of phosphorylated Cdc28 **(inactive form)**, which in turn can lead to G2 delay/stress and therefore many gene expression changes.

**S4 Fig. Hierarchical clustering of positive and negative genetic interaction GSTF pairs.**

Hierarchical clustering of 44 GSTF pairs according to their genetic interaction effects after slow growth correction. These pairs include both negative and positive genetic interactions. Layout and analysis similar to Fig 2.

**S5 Fig. Provided tokens to places in WT condition and transition firing rules**

(A) Provided tokens to regulators depending on edges between them. (B) Transition firing rules for activation and inhibition edges depending on the presence of tokens in upstream places.

